# Role of bradykinin type 2 receptors in human sweat secretion: translational evidence does not support a functional relationship

**DOI:** 10.1101/2020.12.01.407114

**Authors:** Thad E. Wilson, Seetharam Narra, Kristen Metzler-Wilson, Artur Schneider, Kelsey A. Bullens, Ian S. Holt

**Affiliations:** Division of Biomedical Sciences, Marian University College of Osteopathic Medicine, Indianapolis, IN, USA; Department of Physiology, University of Kentucky College of Medicine, Lexington, KY USA; Department of Physical Therapy; Indiana University, Indianapolis, IN USA; Department of Anatomy, Cell Biology, & Physiology; Indiana University, Indianapolis, IN USA; Department of Dermatology; Indiana University, Indianapolis, IN USA

**Keywords:** intradermal microdialysis, HOE-140, bradykinin type 2 receptors, kallikrein-kinin system

## Abstract

Bradykinin increases skin blood flow via a cGMP mechanism but its role in sweating *in vivo* is unclear. There is a current need to translate cell culture and non-human paw pad studies into *in vivo* human preparations to test for therapeutic viability for disorders affecting sweat glands. Protocol 1: physiological sweating was induced in 10 healthy subjects via perfusing warm (46-48°C) water through a tube-lined suit while bradykinin type 2 receptor (B2R) antagonist (HOE-140; 40 μM) and only the vehicle (lactated Ringer’s) were perfused intradermally via microdialysis. Heat stress increased sweat rate (HOE-140 = +0.79±0.12 and vehicle = +0.64±0.10 mg/cm^2^/min), but no differences were noted with B2R antagonism. Protocol 2: pharmacological sweating was induced in 6 healthy subjects via intradermally perfusing pilocarpine (1.67 mg/ml) followed by the same B2R antagonist approach. Pilocarpine increased sweating (HOE-140 = +0.38±0.16 and vehicle = +0.32±0.12 mg/cm^2^/min); again no differences were observed with B2R antagonism. Lastly, 5 additional subjects were recruited for various control experiments which identified that a functional dose of HOE-140 was utilized and it was not sudorific during normothermic conditions. These data indicate B2R antagonists do not modulate physiologically-or pharmacologically-induced eccrine secretion volumes. Thus, B2R agonist/antagonist development as a potential therapeutic target for hypo- and hyperhidrosis appears unwarranted.

## Introduction

Evaporative cooling is vital for human thermoregulation and is primarily achieved via activation of eccrine sweat glands in non-glabrous (thin) skin [1]. Additionally, sweat secretions aid in skin hydration and control of flora and participate in immune defense by the skin [2]. Eccrine sweat glands consist of a bulbous coil which secretes a precursor fluid into the gland lumen; this fluid is then modified in the duct prior to the secretion of sweat onto the skin surface. The secretory clear cells of the coil are under sympathetic nervous system control [3, 4], responding to both thermal and non-thermal factors [5, 6], and have been implicated in both anhidrosis and hyperhidrosis [7, 8]. Despite this nervous system control, the gland is also regulated by local interstitial factors such as osmolality and ion concentration [9, 10]. Thus, it is possible that regulatory compounds could either have direct effects on the gland or may alter local interstitial fluid and ion properties to affect eccrine secretions.

Bradykinin is a dermal kallikrein-kinin system product [11] that is elevated in the plasma of individuals with contact dermatitis, atopic dermatitis and psoriasis [12]. Individuals with these skin conditions can present with altered sweat function, most notably in atopic dermatitis [13, 14]. Bradykinin can induce a wheal and flare response [15] and increases skin blood flow via a cGMP mechanism but does not mediate active cutaneous vasodilation associated with cholinergic stimulation [16]. Eccrine sweat gland activation stimulates bradykinin-forming enzymes and bradykinin formation in the local milieu [17] and in secretions [18]. The presence of glandular kallikrein cells in eccrine sweat glands [19] suggests involvement of the kallikrein-kinin system in ion transport. Lysylbradykinin has been identified as a potential agonist via its ability to mediate short-circuit currents in cultured human sweat gland cells [20, 21]. Thus, given the potential for both direct and indirect effects of bradykinin on eccrine sweat glands, we tested the central hypothesis that bradykinin-induced eccrine secretion rates could be translated into an *in vivo* human preparation.

## Materials and Methods

Study participants (n=18) were healthy; hydrated (urine specific gravity <1.030); normotensive (<140/90 mmHg); non-smokers; not severely obese (BMI <35 kg/m^2^); and with no co-morbidities as determined by a health history and physical exam that included a 12-lead ECG. All participants provided informed written consent for this Marian University Biomedical Sciences IRB approved study that adhered to Declaration of Helsinki guidelines.

*Protocol 1 (Heat Stress):* Ten participants had 2 intradermal 30KDa microdialysis membranes (BASi, West Lafayette, IN, USA) inserted into the dorsal forearm; membranes were perfused with 40 μM HOE-140 (Sigma-Aldrich, St. Louis, MO, USA), a bradykinin type 2 receptor (B2R) antagonist [22], and lactated Ringer’s vehicle (Baxter, Deerfield, OH, USA). To induce physiological sweating, 46-48°C water was perfused through a high-density tube-lined suit until intestinal pill (HQ Inc., Palmetto, FL, USA) temperature increased by 1°C. Intestinal pill temperature was used as an index of internal temperature [23]. Surface sweat rate was obtained just superficial to the membrane via capacitance hygrometry. Cutaneous vascular conductance (CVC) was measured via laser Doppler flowmetry (Moor Instruments, Wilmington, DE, USA) and arterial blood pressure (Finapres Medical Systems, Enschede, Netherlands; GE Healthcare, Boston, MA, USA). Additionally, after sustained sweating and internal temperature measurements were obtained, both microdialysis membranes were perfused with bradykinin (1 mM; Sigma-Aldrich) to assess B2R inhibition and uncover potential bradykinin involvement at the control site (Figure 1A). *Protocol 2 (Pilocarpine):* Six participants had 2 intradermal microdialysis membranes inserted and perfused with 1.67 mg/ml pilocarpine (Sigma-Aldrich) prior to the same B2R antagonist and agonist approach described in *Protocol 1* (Figure 1B). Drug concentration used in protocols mimicked previously published intradermal microdialysis studies [16, 24]. Five additional participants were recruited for various control experiments to: 1) identify potential independent effects of HOE-140 on sweating (Figure 1C), 2) determine if there were additive effects of bradykinin and pilocarpine on CVC (Figure 1A-B), and 3) verify functional cutaneous B2R antagonism via CVC and protein extravascularization (BCA protein assay of dialysate; Thermo Scientific, Waltham, MA, USA; Figure 1D). Increases in skin flux (vasodilation) and dialysate protein concentration (greater capillary permeability) have previously been identified during bradykinin administration in the skin [25, 26].

**Figure 1.**
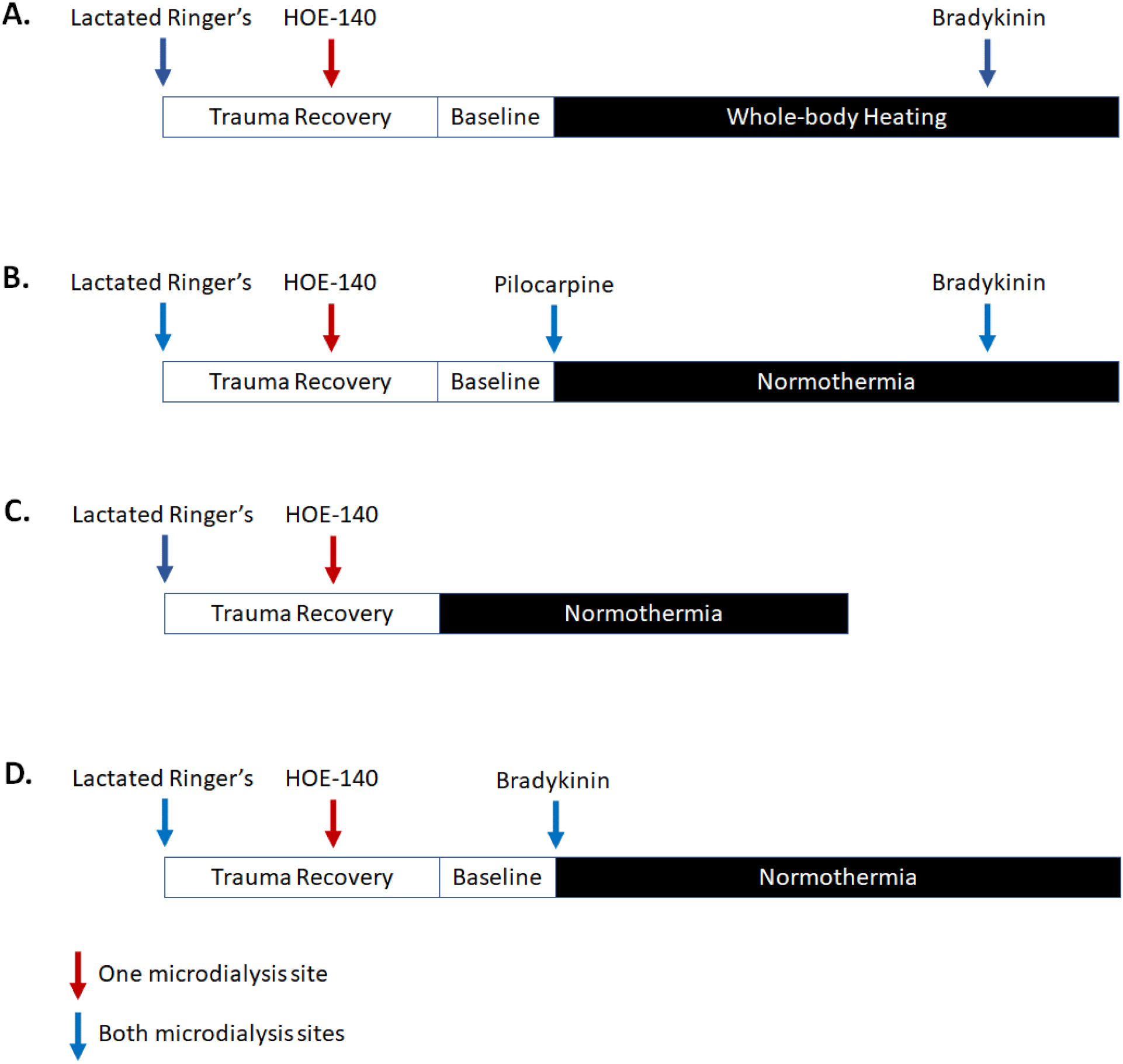
Experimental schematic for Protocol #1 (panel A), Protocol #2 (panel B), and additional control and development experiments (panels C-D).

Data were collected at 100 Hz frequency via an analog to digital converter (BioPac Inc., Santa Barbara, CA, USA). Sweat rate was defined as the volume secreted onto the skin surface per unit time above basal transepithelial water loss (TEWL). Measurements that were collected across time were analyzed between group (HOE-140 vs. vehicle) via repeated measures ANOVA. Pre-post measurements and single point drug comparisons were analyzed via dependent and independent T-tests, respectively. Onset of sweating and slope of the relationship between the increase in sweat rate and internal temperature were determined graphically by a single blinded investigator [27]. Statistical significance was set at p<0.05.

## Results

### Protocol 1

Whole-body heat stress increased internal (37.1±0.1 to 37.9±0.1°C) and uncovered skin (30.1±0.4 to 32.9±0.4°C) temperatures, resulting in increased sweat rate (Figure 2A) and CVC (63±11 to 181±22 and 85±15 to 204±19 flux/mmHg for HOE-140 and vehicle, respectively). HOE-140 and vehicle sites were not different for either sweat rate or CVC, indicating a lack of a B2R antagonist effect. Heat stress increased heart rate (62±3 to 94±6 bpm) and systolic blood pressure (112±3 to 121±5 mmHg), decreased diastolic blood pressure (66±2 to 58±2 mmHg), and did not alter mean arterial blood pressure (83±2 to 82±2 mmHg). No group differences were observed in the temperature threshold for the onset of sweating between HOE-140 (0.15±0.06) and vehicle (0.19±0.07°C). Similarly, the slope of the relationship between the increase in sweat rate and internal temperature was not different between the two sites (HOE-140 = 1.14±0.17 and vehicle = 0.98±0.15 mg/cm^2^/min/°C Δ in internal temperature). The addition of exogenous bradykinin (1 mM) in the perfusate did not modulate sweat rate during whole-body heating in either HOE-140 or vehicle sites.

**Figure 2.**
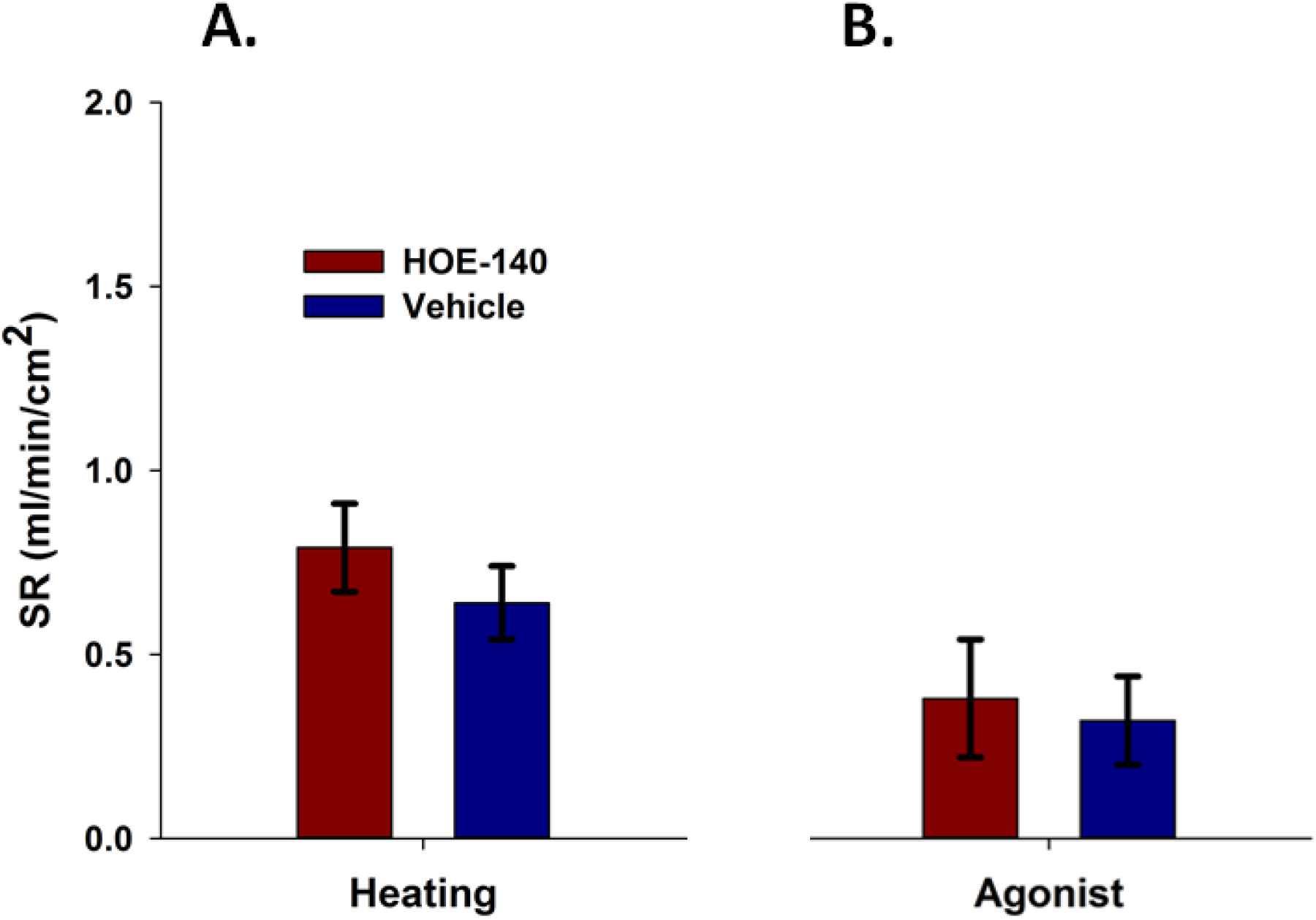
Effect of bradykinin type 2 receptor antagonism (HOE-140) on heat-induced (panel A; n=10) and cholinergic agonist (pilocarpine)-induced (panel B; n=6) sweating in human forearm skin. SR denotes sweat rate. No HOE-140 vs. vehicle differences were noted in either panel A or B. Data are presented as mean ± SEM.

### Protocol 2

Pilocarpine induced sweating (Figure 2B) and increased CVC (88±36 to 183±55 and 73±25 to 208±66 flux/mmHg for HOE-140 and vehicle, respectively) but again, no differences between sites were noted with either variable. The addition of exogenous bradykinin (1 mM) in the perfusate also did not modulate sweat rate during pilocarpine perfusion (*Protocol 2*) in either HOE-140 or vehicle sites.

### Additional Control Experiments

Because of the novel drug perturbation we performed several pilot and control studies to identify potential collateral effects, use of functional dose, and potential additive effects. HOE-140 *per se* delivered during normothermia was also identified not to be sudorific (i.e., no water loss above TEWL), indicating that HOE-140 does not affect sweating independently. 40 μM HOE-140 attenuated the 1 mM bradykinin-induced cutaneous vasodilation and protein extravascularization, indicating a functional antagonist dose was employed. When 1 mM bradykinin was combined with 1.67 mg/ml pilocarpine, no further increases in skin blood flow were observed compared to pilocarpine alone.

## Discussion

Although the kallikrein-kinin system is present in eccrine sweat glands [11, 19], these data indicate that B2R antagonists do not modulate the absolute rate of physiologically-or pharmacologically-induced sweating. Similar to cholinergic stimulation, B2R activation increases cytosolic Ca^2+^. Eccrine sweat glands are dependent on Ca^2+^ for cholinergic secretion; attenuating interstitial-to-cytosolic Ca^2+^ gradients and entry decreases or abates sweating in cell culture and *in vivo* [28]. B2R activation also increases cytosolic Ca^2+^ and is associated with reorganizing cytoskeleton F-actin, leading to changes in transepithelial resistance [29] and redistribution of occludin tight junctions, which increases protein diffusion in various epithelial tissues [30]. These actions could have, in theory, increased sweat secretions, but this was not our observation using our experimental setup.

Brenglemann et al. [31] identified a lack of cutaneous vasodilation in patients with anhidrotic ectodermal dysplasia, suggesting a mechanistic link between sweating and blood flow. Previous studies have demonstrated an association between bradykinin formation and periglandular vasodilation in human forearm skin [32], but not indicated the mediator compound of active vasodilation [16]. The bradykinin-induced blood flow to secretion rate relationship appears to hold in related exocrine glands, i.e., salivary glands [33, 34], but neither our B2R agonist nor antagonist data support this link in eccrine sweat glands.

*In vivo* delivery of compounds via intradermal microdialysis is a translational methodological advancement over previous techniques and provides useful data to translate findings from isolated sweat glands, cell line (NCL-SG3), and cell culture [28, 35]. To improve generalizability and robustness of the data and interpretations, we utilized a dual *in vivo* approach to engage sweating: a physiological (heat stress) approach for presynaptic acetylcholine release and a pharmacological (pilocarpine) approach to directly bind postsynaptic M3 receptors. It is possible that greater doses of bradykinin could have yielded different responses. However, because bradykinin is a nociceptor agonist, *in vivo* investigations are somewhat limited in terms of maximal delivery dose to unanesthetized participants.

In conclusion, B2R may have condition-specific or supportive roles for epithelial transport, but our current *in vivo* data do not support a translational bradykinin mechanism related to absolute sweat output. Thus, B2R agonist/antagonist development as a potential therapeutic target for hypo- and hyperhidrosis appears unjustified at this time.

## Acknowledgements

Additional technical assistance was provided by Malavika Seetha. Additional logistic support was provided by the Hill-Rom Simulation Center at Marian University.

## Funding Sources

Funding was provided by an external grant from National Institute of Arthritis, Musculoskeletal, and Skin Diseases [AR-069912] to TEW and KMW, and the following internal sources: Faculty Research Development grant to TEW and the summer DO research fellowship program from Marian University College of Osteopathic Medicine to AS, KB, and ISH.

## Notes

### Competing Interest Statement

The authors have declared no competing interest.

